# Tetanus Toxoid Utilizes Dual-Faceted Anticancer Mechanism Through Targeting Tumoral Sialic Acids and Enhancing Cytotoxic CD4+ T cell Responses Against Pancreatic Cancer

**DOI:** 10.1101/2024.11.19.624337

**Authors:** Eileena F Giurini, Sam G Pappas, Kajal H Gupta

**Affiliations:** Division of Surgical Oncology, Department of Surgery, Rush University Medical Center, Chicago, IL, USA 60612; Division of Pediatric Surgery, Department of Surgery, Rush University Medical Center, Chicago, IL, USA 60612

## Abstract

Pancreatic ductal adenocarcinoma (PDAC) remains profoundly resistant to conventional chemotherapy and immunotherapeutic interventions. Innovative therapeutic modalities, particularly microbe-derived immunotherapies, have demonstrated durable anti-tumor efficacy in preclinical PDAC models. This study elucidates that administration of the FDA-approved Haemophilus influenzae type b (H Flu - Hiberix) vaccine attenuates tumor progression and enhances survival outcomes in murine PDAC. H Flu treatment significantly augmented CD4+ T cell, CD8+ T cell, and natural killer (NK) cell infiltration within the tumor microenvironment, concurrently inducing a cytotoxic T cell phenotype, evidenced by upregulation of CD69, granzyme B, and perforin. Additionally, H Flu therapy promoted the accumulation of CD44+ CD62L-memory T cells within tumors of pre-immunized mice. Mechanistic investigations revealed that depletion of CD4+ T cells or NK cells, but not CD8+ T cells, negated the anti-tumor efficacy of H Flu, suggesting that CD4+ T cells and NK cells are critical mediators of H Flu-induced anti-tumor immunity. To further elucidate the mechanistic basis of H Flu’s anti-tumor activity, we assessed the individual constituents of the H Flu vaccine: tetanus toxoid (TT) and polyrobosyl ribitol phosphate (PRP). Notably, TT administration achieved superior tumor growth suppression, characterized by enhanced CD4+ T cell cytotoxicity and increased NK cell infiltration, relative to PRP or PBS-treated controls. Furthermore, TT induced apoptosis in PDAC cells and reduced their proliferation, potentially by targeting tumor-associated sialic acids. This disruption might interfere with the interaction between sialic acids and siglec receptors, thereby impairing mechanisms of immune evasion.TT-mediated modulation of sialic acid expression in cancer cells underscores its potential to augment immunotherapeutic efficacy in PDAC. Collectively, these findings reveal a novel anti-cancer mechanism for TT, leveraging both immunostimulatory and sialic acid-targeting pathways to suppress PDAC progression.

**Graphical Abstract:** 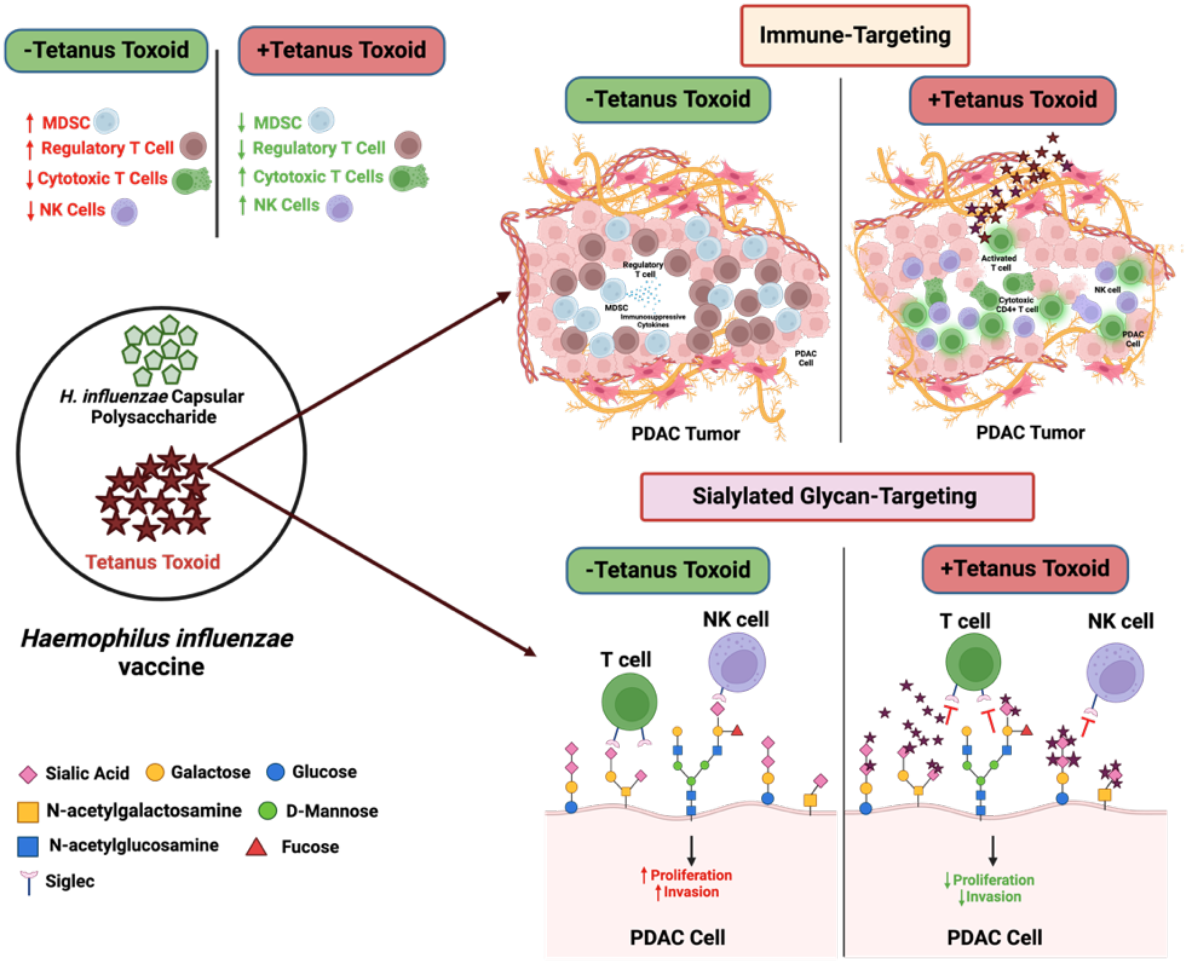

## Introduction

Pancreatic ductal adenocarcinoma (PDAC) is a devastating malignancy responsible for nearly 5% of all cancer deaths worldwide^1^. Despite the increasing incidence and aggressive nature of PDAC, little progress has been made in improving the 5-year survival rate of 12%, which has remained stagnant for over 20 years^2^. These poor outcomes can largely be attributed to the challenges in treatment of PDAC. The advent of immune checkpoint inhibitor (ICI) therapies in cancer care has culminated in dramatic improvements for many solid tumor types, however, these same therapies are effective in <1% of PDAC patients^3,4^. The resounding failure of ICI therapies in PDAC can be ascribed to poor tumor immunogenicity, limited neoantigen generation, and a fibrotic stromal barrier surrounding the tumor. Importantly, the stromal layer precludes entry to effector cells as well as therapeutic agents, while sustaining a highly immunosuppressive tumor microenvironment (TME) that is detrimental to immune activation^5^.

While most clinically utilized cancer immunotherapies solely target CD8+ T cell-mediated tumor eradication^6^, there is emerging evidence that CD4+ T cell and natural killer (NK) cell populations also have great potential as a therapeutic approach^7,8^. In the context of cancer, CD4+ T cells exert antitumor responses through two primary mechanisms: indirect and direct cytotoxicity. Canonically, CD4+ T cells perform a dynamic helper role, promoting priming, attenuating negative regulation, and supporting establishment of immunological memory for cytotoxic CD8+ T cells^7,9^. In contrast, direct CD4+ T cell cytotoxicity occurs through secretion of cytolytic granzymes or effector cytokine release following antigen presentation from major histocompatibility class II (MHCII) complexes^10,11^. Given their unique tumor recognition mechanism, NK cell engagement is also a promising immunotherapy strategy. NK cells initiate antitumor immunity independently of antigen presentation from MHC, rather through recognition of “stress” ligands uniquely expressed by tumor cells^8^. Because NK cells possess the ability to initiate antitumor immunity without antigen presentation, they are suitable for use against tumors with limited antigenicity. NK cells typically mitigate tumor progression through release of cytotoxic mediators, perforin or granzymes, as well as inducing antibody-dependent cell mediated cytotoxicity (ADCC) in tumor cells through CD16A activation^12^. Thus, directed engagement of NK cells or CD4+ T cells can ultimately culminate in activation of multiple effector cell populations for a potentiated antitumor response.

Given that many solid tumors acquire resistance to conventional and immunotherapies over time, significant efforts have been directed into investigating microbial based cancer therapies (MBCTs) as a novel therapeutic intervention^13^. MBCTs can provide tumor-selective delivery of cytotoxic agents, and often possess intrinsic oncolytic abilities^13^. Importantly, MBCTs are also able to introduce “non-self” microbial antigens into the tumor, enhancing recognition by the host immune system^13,14^. Following initial host recognition, MBCTs often trigger effector programs in lymphocyte populations, culminating in T cell or NK cell-mediated antitumor immunity^15^. MBCT induced T cell activation can occur through enhancing antigen presentation and co-stimulation from antigen presenting cells (APCs)^14^. Alternatively, MBCTs can directly engage immune sensors expressed on T cells or NK cells, that when activated, trigger a pro-inflammatory signaling cascade^14,16,17^. Even for highly immunosuppressive tumors such as PDAC, MBCTs have demonstrated robust therapeutic efficacy. Recently, delivery of immunostimulatory antigens by *Listeria monocytogenes* and *Salmonella typhimurium* to PDAC tumors potently activated cytotoxic T cell populations, leading to durable reduction in primary tumor burden and metastasis^18,19^.

Live bacteria or active viral MBCTs have promising anticancer effects, however, their usage can culminate in severe adverse toxicities^13^. As an alternative to live-bacteria MBCTs, our group has investigated repurposing microbial-based vaccines as cancer immunotherapies^20-23^. Microbial-based vaccines provide much of the same strong microbial antigens present in live MBCTs, but in a less toxic formulation. Using this principle, we demonstrated that intratumoral administration of the seasonal influenza vaccine or inactivated SARS-CoV-2 vaccine culminated in reduced tumor progression^20,22^. The physiological response was accompanied by increased cytotoxic T cell infiltration into the tumor, indicating the introduction of microbial antigens into the tumor rendered it immunologically “hot” ^20-22^. In a subsequent study, we identified that the interaction between influenza vaccine derived hemagglutinin and tumoral sialic acids greatly contributed to the antitumor effect from the vaccine, indeed highlighting sialic acids as a potential therapeutic focus for vaccine based MBCTs^23^.

In the context of cancer, aberrant glycan sialylation of cancer cells promotes tumorigenesis through tumor extrinsic and intrinsic mechanisms^24^. Accordingly, abnormal sialylation has been correlated with a poor prognosis for colorectal, prostate, and pancreatic malignancies^25-27^. Expression of sialic acids on the surface of tumor cells evades immune recognition predominantly through interacting with sialic acid binding immunoglobulin type lectins (Siglecs). On many immune cell types, including T cells and NK cells, sialic acid binding with Siglecs initiate an inhibitory program within the cell. This interaction diminishes activation and cytotoxicity of these cells, consequently protecting tumor cells from eradication^28,29^. It is this interaction that the Siglec-sialic acid axis is now recognized as a novel immunotherapy checkpoint^30^. Sialylation also contributes to many functional abilities acquired by tumor cells. Indeed, hypersialylation of membrane proteins and integrins, effectively inhibits apoptosis and other cell death related intracellular signaling^24,29^. Additionally, sialylation of cell surface adhesion molecules has been implicated in promoting invasion and metastasis, augmenting the aggressiveness of the disease^29^. The potential for enhancing antitumor immunity and mitigating pro-tumoral signaling with a singular target, position sialylation targeting as a favorable therapeutic approach.

In this study, we build upon our established vaccine-based cancer treatment paradigm, investigating the therapeutic potential of clinically utilized microbial based vaccines as an immunotherapy for PDAC. We report that the FDA-approved *Haemophilus influenzae* serotype B (H Flu) vaccine reduces tumor progression and improves survival in subcutaneous, orthotopic, and genetically engineered murine models (GEMM) of PDAC when administered intratumorally or intraperitoneally. Accordingly, we elucidated integral immunological mechanisms mediating antitumor immunity, and the contribution of each vaccine component to the antitumor properties of H Flu. The Haemophilus influenzae (H flu) vaccine consists of two key components: polyribosylribitol phosphate (PRP) and tetanus toxoid (TT). Through this investigation we identify the tetanus toxoid component as the predominant contributor of the tumor immunogenicity of H Flu, as well as interacting with sialic acids expressed on PDAC tumor cells. We further identify that the tetanus toxoid and sialic acid interaction possesses antiproliferative and anti-invasive effects on tumor cells. In conclusion, this study describes tetanus toxoid as a dual-faceted immune and tumoral sialic acid-targeting therapy against PDAC.

## Materials and Methods

### Mice

C57BL/6 mice aged 6 to 8 weeks, used to generate the 6419c5 (subcutaneous) and KPC-luc (orthotropic) tumor models, were purchased from The Jackson Laboratory. Pdx1-Cre × LSL-*Kras*^G12D^ × LSL-*TP53*^R172H^ (KPC) mice were kindly provided by Dr. Ajay Rana (University of Illinois at Chicago). Male mice were used for all experiments conducted with C57BL/6 mice, and female mice were used for all experiments with KPC mice. Animals were housed in specific-pathogen-free facilities and all experimental procedures were in accordance with policies established by the Institutional Animal Care and Use Committee (IACUC) at Rush University Medical Center.

### Cell Lines

The 6419c5 cell line was isolated from a late-stage primary PDAC tumor in a KPCY mouse and was purchased from Kerafast^31^. The KPC-luc cell line was generated from a tumor in a Pdx1-Cre × LSL-*Kras*^G12D^ × LSL-*TP53*^R172H^ mouse followed by luciferase gene insertion into the cell line through lentiviral transduction^32^. The KPC-luc cell line was kindly provided by Dr. Bassel El-Rayes (University of Alabama-Birmingham). Both cell lines were cultured using DMEM supplemented with 2 mM glutamine, 10% FBS, 100 units/mL penicillin, and 100 mg/mL streptomycin. The HEK-Blue TLR reporter cell lines were stably transfected with plasmids containing a single murine TLR gene and a NF-kB inducible secreted embryonic alkaline phosphatase (SEAP) gene to measure TLR stimulation. The expression of TLR and SEAP plasmids was maintained through supplementing the aforementioned complete DMEM medium with 30 µg/mL blasticidin and 100 µg/mL zeocin as selection antibiotics. Measurement of TLR activation was assessed as previously described^20,21^.

### Tumor Development

For experiments using the 6419c5 tumor model, 5 × 10^5^ cells suspended in PBS were subcutaneously administered into the right flank of C57BL/6 mice. Changes in tumor growth were monitored with caliper measurements in two perpendicular experiments. For the KPC-luc orthotopic model, C57BL/6 mice were initially anesthetized with a ketamine/xylazine cocktail (90 mg/kg and 10 mg/kg, respectively). Anesthesia was maintained through isoflurane inhalation, and animals were placed on their right side, exposing the area of the spleen in which the hair surrounding the area had been removed the day prior. Povidone iodine solution followed by 70% isopropyl alcohol was used to sterilize the area prior to incision. A 1 cm incision was made through the skin and musculature, exposing the underlying organs. The pancreas and spleen were gently pulled out of the incision site using a sterile cotton tipped applicator. Once situated, 8 × 10^4^ KPC-luc cells in a 30% Matrigel solution were injected into the tail of the pancreas. After injection, the pancreas and spleen were returned to their correct anatomical position and musculature and skin were sutured using 6-0 absorbable and 3-0 non-absorbable sutures, respectively, to close the incision. Changes in tumor growth was monitored through intraperitoneal injection of 150 mg/kg D-luciferin, followed by bioluminescence measurement using an in vivo imaging system (IVIS), once or twice weekly.

### Survival Studies

Following completion of the treatment regimen, KPC and KPC-luc tumor bearing mice were monitored and weighed for indications of decline in health, once or twice a week. Monitoring was continued until animals had succumbed to disease, or had met health-related endpoints requiring euthanasia, as per IACUC policies.

### *In Vivo* Depletion Studies

*In vivo* depletion of CD4+ T, CD8+ T, and NK cells were conducted using anti-CD4 (clone GK1.5), anti-CD8 (clone YTS169.4), and anti-NK1.1 (clone PK136) purchased from BioxCell. 300 µg of anti-CD4, anti-CD8, anti-NK1.1 was administered to KPC-luc tumor-bearing mice via intraperitoneal injection, twice or once weekly, according to treatment protocol. The respective isotype for each depletion antibody was administered to control groups.

### Vaccine and Vaccine Components

The FDA-approved Hiberix vaccine indicated for immunization against *Haemophilus influenzae* serotype B is manufactured by GlaxoSmithKline. The vaccine was obtained for this study through Rush University Medical Center Pharmacy. H Flu was reconstituted with PBS as described in the FDA package insert. Once reconstituted, 50 µl of undiluted vaccine was administered intratumorally for subcutaneous model. For orthotropic model intraperitoneal administration 100 µl of vaccine was used. For the mechanistic component studies, the formaldehyde-inactivated tetanus toxoid (TT) derived from *Clostridium tetani* was purchased from Sigma Aldrich (#582231) and Enzo Life Sciences (#ALX-630-108-C100). The Haemophilus b capsular polysaccharide, polyribosyl-ribitol-phosphate (PRP), was purchased from Creative Diagnostics (#DALG0481). Both components were reconstituted in PBS to deliver a 1 µg dose of PRP or 2.5 µg TT was administered via intraperitoneal injection, the same amount found in a 100 µl dose of H Flu. PBS was used as a control for all vaccine and component studies.

### Flow Cytometry

Tumors were processed to get single cell suspension as previously described^20^. To identify viable cells, Live/Dead Fixable Aqua Dead Cell Stain Kit (405 nm excitation) was added to antibody staining cocktails. To identify immune cells in the tumor samples, anti-CD45 antibodies were used. Using the CD45+ population, T cells were identified using a CD3-targeting antibody, CD8+ T cells were identified using CD3 and CD8-specific antibodies, and CD4+ T cells were identified using CD3 and CD4 specific antibodies. Natural killer cell populations were identified using NK1.1 and CD3 (negative gating) antibodies. Myeloid derived suppressor cells were identified using CD11b and Gr-1 specific antibodies. Extracellular staining cocktails also included antibodies specific to CD62L, CD44, LAG-3, and CD69. The amount of each antibody used for extracellular staining was determined by the manufacturer’s instruction. For intracellular staining, samples were permeabilized and fixed using Cytofix/Cytoperm per the manufacturer’s instructions prior to staining with FOXP3, Granzyme B, and perforin targeting antibodies. The respective isotype controls were run in simultaneous with antibody-stained samples. Sialic acid expression was determined using biotinylated SNA (α-2,6 sialic acid) and MALII (α-2,6 sialic acid) lectins purchased from Vector Laboratories. 1 × 10^5^ 6419c5 cells were incubated with 2 µg/mL of biotinylated lectin for 45 minutes at 4**°**C. Detection of lectins was completed using 1 µg/mL of streptavidin conjugated AlexaFluor 488 (Invitrogen), followed by a 20 minute incubation at RT. All flow cytometry was completed using the BD LSRFortessa. Flow cytometry analysis was then analyzed using FlowJo (version 10.10).

### Histology

Resected tumor samples were placed in 10% formalin for 24 hours followed by 70% ethanol for at least 24 hours. Samples were then dehydrated with an increasing ethanol gradient and embedded in paraffin blocks. 5 µm sections were cut from the blocks using a microtome and placed on slides. Deparaffinization was completed through two xylene washes and rehydrated with a decreasing ethanol gradient. Hematoxylin and eosin staining was completed using an H&E staining kit purchased from Abcam, following the manufacturer’s instructions. Stained sections were then imaged using a Keyence BZ-X810 microscope.

### Tetanus Toxoid Immunoprecipitation

6419c5 cells were cultured to 80% confluency, then treated with 1.5 µg/mL TT, H Flu (also containing 1.5 µg/mL TT), or PBS for 1 hour at 37**°**C. Following treatment, cells were washed with ice cold 1X PBS then lysed with ice cold cell lysis buffer for 30 minutes on ice. A HEPES-based cell lysis buffer used in these experiments, adopted from a previously described recipe^33^. Following incubation, samples were centrifuged at 16000 x g for 15 minutes at 4**°**C and supernatant was collected. Total protein was subsequently quantified using a Micro BCA Protein Assay Kit (Thermo Scientific). In simultaneous, Dynabeads (Thermo Scientific) coupled to an anti-tetanus toxin antibody, using 60 µg of antibody per sample. Appropriate incubation, blocking, and wash steps were completed, following manufacturer’s instruction. Antibody-coupled Dynabeads were then combined with 1.75 mg total cell protein and incubated for 24 hours at 4**°**C, with constant rotating. The subsequent washing and protein elution steps for each sample was completed following an established STAR Protocol^33^. To confirm TT interaction with PDAC cells, the immunoprecipitates were combined with an appropriate volume of 6X Laemmli buffer and heated for 5 minutes at 95**°**C. Samples were subsequently subjected to SDS-PAGE separation. After SDS-PAGE, the protein gel was stained using GelCode Blue Safe Stain (Thermo Scientific) and destained with ultrapure water, according to manufacturer’s instruction. Once the protein gel was destained, the presence of TT was visualized using a Licor Odyssey M imaging system (LICORbio).

### Sialic Acid Binding Assay

5 × 10^4^ 6419c5 cells were seeded in opaque black 96-well plates, previously coated with 50 µg/mL poly-D-lysine solution. The following day, cells were treated with 150 µM of sialyltransferase inhibitor P-3Fax-Neu5Ac (R&D Systems, #5760/10) in culture medium or a vehicle control, as previously described^23^. After incubation, cells were treated with H Flu, TT, PRP, or a PBS control for 1 hour at 37**°**C. Cells were then fixed using 4% formaldehyde solution for 15 minutes at RT and subsequently washed with PBS. Following fixation, cells were blocked using a 5% bovine serum albumin solution for 1 hour. Bound TT was identified using a fluorescent conjugated anti-tetanus toxin antibody (Novus Biologicals) at a 5 µg/mL concentration, and was incubated for 2 hours at RT. Cells were then washed thrice with PBS, and fluorescent intensity (bound TT) was quantified using the Varioskan Lux (Thermo Fisher).

### Cell Proliferation Assay

1 × 10^5^ 6419c5 cells were seeded in 6-well plates. The following day, the cells were treated with 1.5 µg/mL TT combined with an appropriate vehicle control or 1.5 µg/mL SNA lectin (Vector Laboratories), MALII lectin (Vector Laboratories), or N-acetylneuraminic acid (Neu5Ac, Sigma Aldrich) in culture medium. Cells were incubated with the aforementioned treatments for 48 hours at 37**°**C. Following incubation, changes in cell proliferation were determined using the Click-iT EdU Alexa Fluor 488 Flow Cytometry Kit (Thermo Fisher), following the manufacturer’s instruction.

### Annexin V Apoptosis Assay

6419c5 cells were seeded in opaque black 96-well plates at a density of 5 × 10^4^ cells per well. The following day, the RealTime-Glo Annexin V Apoptosis Kit (Promega) was according to manufacturer’s instruction to detect apoptosis in real time. Cells were incubated with 1.5 µg/mL TT combined with 1.5 µg/mL SNA lectin (Vector Laboratories), MALII lectin (Vector Laboratories), or Neu5Ac (Sigma Aldrich) or an appropriate vehicle control for 48-72 hours at 37**°**C. Annexin V was measured at 48- and 72-hour timepoints using the Varioskan Lux (Thermo Fisher).

### Statistical Analysis

Statistical analyses were completed with GraphPad Prism Version 10.1.1. A two-way ANOVA with Tukey correction was used to compare data with multiple timepoints. For a single comparison between two groups, a paired Students’ t-test was used. For comparison between more than two groups, a 1-way ANOVA was used. To ascertain statistical significance in survival studies, a Log-rank (Mantel-Cox) curve was generated. P values < 0.05 were considered statistically significant. *P < 0.05, **P < 0.01, ***P < 0.001, and ****P < 0.0001.

## Results

### Systemic administration of H Flu vaccine elicits antitumor immunity in subcutaneous PDAC model

Asplenic vaccines are microbial-based vaccines often administered to PDAC patients prior to splenectomy, as a means to reduce sepsis^34^. Building on our group’s expertise in repurposing microbial-based vaccines for cancer immunotherapy, we hypothesized that asplenic vaccines could also serve as effective immunotherapies for pancreatic ductal adenocarcinoma (PDAC). To explore their therapeutic potential, we conducted an initial screen of asplenic vaccines and identified *Haemophilus influenzae* serotype B (H Flu) vaccine as a promising candidate. We challenged C57BL/6 mice with the 6419c5 cell line, derived from a KPCY mouse tumor which generates tumors with a characteristic immunosuppressive TME^35^. Established tumors were treated with H Flu or a PBS control via intratumoral injection (Supplemental Figure S1A). Notably, tumors treated with H Flu showed a marked reduction in growth compared to PBS-treated controls (Supplemental Figure S1B). Given that direct intratumoral administration is challenging for pancreatic ductal adenocarcinoma (PDAC) due to the need for invasive surgery, we next assessed the antitumor effects of these vaccines via the less invasive, intraperitoneal administration. Using a similar treatment schedule (Supplemental Figure S1C), intraperitoneal administration of H Flu significantly reduced tumor progression compared to PBS. As H Flu demonstrated antitumor activity through both intratumoral and intraperitoneal routes, we focused further investigations on its potential as a therapy for PDAC. An ATP assay measuring viable cells indicated that H Flu treatment decreased the number of metabolically active cancer cells compared to PBS (Supplemental Figure S1E), suggesting that H Flu may possess antiproliferative effects against PDAC cells.

Given the stark contrast between the strong immunogenicity of microbial components and the immunosuppressive tumor microenvironment (TME) of PDAC, we further investigated the effects of H Flu on the intratumor immune landscape. Flow cytometry analysis of H Flu-treated tumors (Supplemental Figure S1F) showed substantial reconfiguration of the TME, with increased infiltration of pro-inflammatory immune cells and a reduction in immunosuppressive cell populations (Supplemental Figure S1G). Specifically, H Flu treatment led to significant increases in CD8+ T cells and natural killer (NK) cells (Supplemental Figure S1H). Moreover, it induced a cytotoxic, effector phenotype in CD4+ T cells, marked by increased granzyme B production (Supplemental Figure S1H). Additionally, H Flu treatment shifted the balance towards a more immune-infiltrated TME by increasing the CD8+ T cell to regulatory T cell (Treg) ratio and reducing the presence of myeloid-derived suppressor cells (MDSCs) within the tumor (Supplemental Figure S1H).

Toll-like receptor (TLRs) are well-recognized for sensing conserved microbial structures, and are also implicated in directing antitumor immunity^21,35^. Because of the microbial origin of the H Flu vaccine, we next investigated its TLR stimulatory potential. Using stably transfected TLR reporter cell lines and their parental (Null) controls, H Flu treatment significantly stimulated TLR2 and TLR4 expressing cells compared to the parental control cells (Supplemental Figure 1I). To elucidate any direct, non-immune mediated, anticancer properties of the H Flu vaccine, we then examined the effect of H Flu on 6419c5 viability. These results indicate that the H Flu vaccine may possess both immunostimulatory and immune-independent anticancer properties in PDAC.

**Figure 1.**
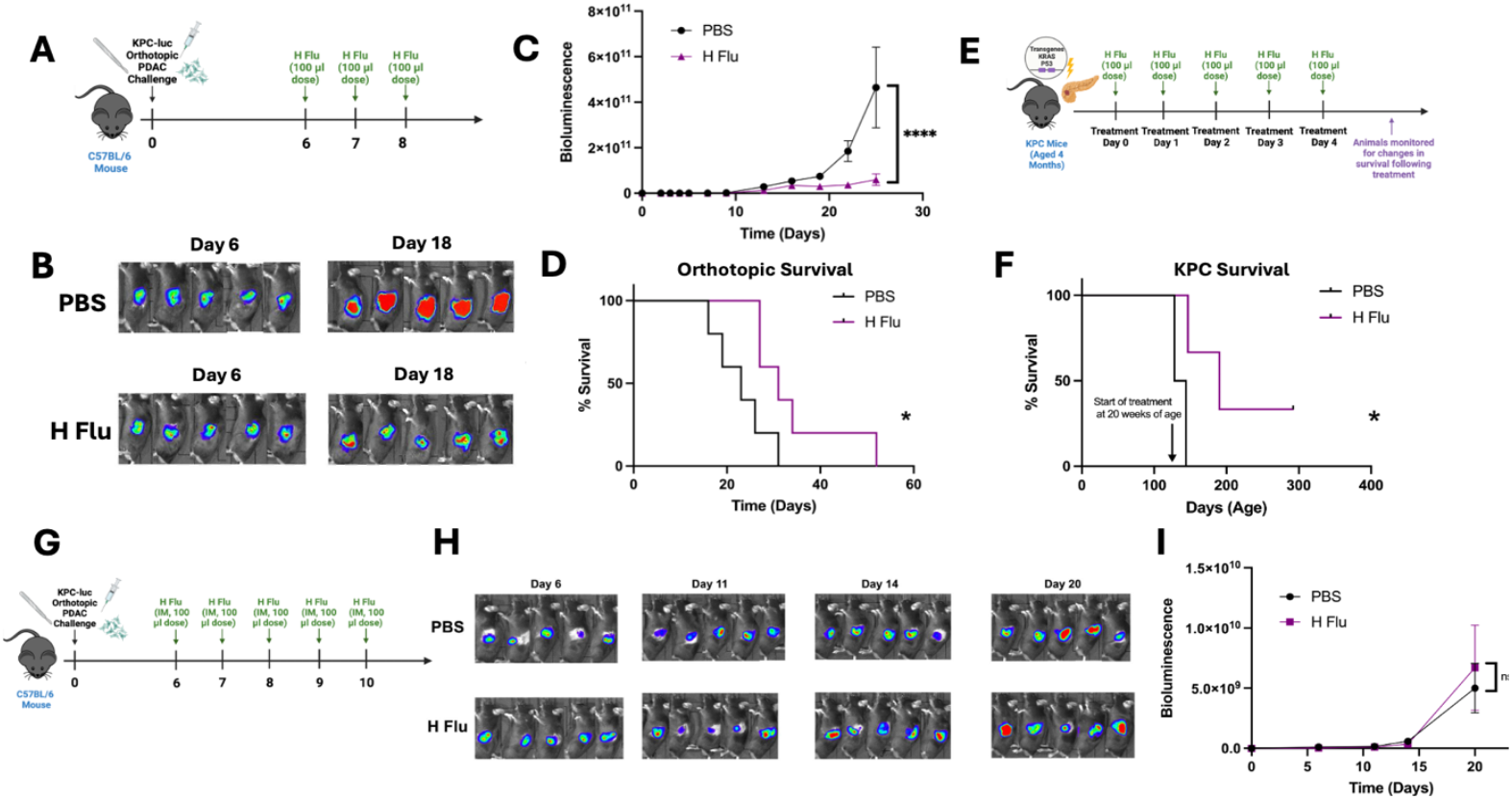
Intraperitoneally Administered H Flu Vaccine Improves Outcomes in Orthotopic and Genetically Engineered PDAC Models. (**A**) Experimental design describing treatment schedule utilizing orthotopic PDAC tumor model. n = 5 mice per group. (**B**) Representative bioluminescent images of mice bearing KPC-luc tumors following H Flu or PBS treatment as described in A. (**C**) Tumor growth curve plotting bioluminescent intensity (p^-1^ sec^-1^ cm^2^ sr^-1^) compared to time following treatment regimen described in A. (**D**) Survival analysis of KPC-luc tumor bearing mice following treatment schedule described in A. (**E**) Experimental design describing treatment schedule of KPC mice. n = 3 mice per group. (**F**) Survival analysis of KPC mice following treatment schedule described in E. (**G**) Experimental design describing intramuscular administration of H Flu in orthotopic PDAC model. n = 5 mice per group. (**H**) Representative bioluminescent images of mice bearing KPC-luc tumors following H Flu or PBS treatment as described in H. (**I**) Tumor growth curve plotting bioluminescent intensity (p^-1^ sec^-1^ cm^2^ sr^-1^) compared to time following treatment regimen described in H. 2-way ANOVA with Sidak’s correction (C and I), Mantel-Cox log rank test (D and F) ns – not significant, *P < 0.05, ****P < 0.0001.

### Intraperitoneally, but not intramuscularly administered, H Flu reduces tumor progression and improves survival orthotopic PDAC tumors

Certainly, the peritumoral environment differs greatly between pancreatic tumors induced in the subcutaneous tissue and those developing in the ducts of pancreas. To more closely emulate the environment surrounding PDAC tumors, we next investigated the therapeutic efficacy of H Flu in an orthotopic and KPC GEM murine mouse model. We utilized a luciferase-tagged cell line, KPC-luc, to observe changes in tumor progression with an in vivo imaging system. Following a similar treatment schedule as in previous experiments (Figure 1A), we observed a drastic reduction in tumor progression in H Flu treated mice, as measured by significantly reduced bioluminescent intensity (Figure 1B & 1C). Consistent with this finding, H Flu treated mice also experienced significant improvements in survival outcomes (Figure 1D). Given that genetically engineered mouse model, KPC mice, are widely regarded as exhibiting many of the hallmarks of human PDAC pathogenesis, we next examined the effect of H Flu utilizing this strain of mice^36^. KPC mice treated with H Flu indeed experienced prolonged survival with a median survival rate of 214 days, compared to a median survival of 136 days in PBS treated mice (Figure 1E & 1F). Further, no drastic weight loss was observed during or after H Flu treatment (data not shown), indicative of the overall health of the animals as well as the tolerability of H Flu as a therapeutic agent.

In clinical settings, the *Haemophilus influenzae* vaccine is intended for intramuscular administration in humans^37^. To determine if intramuscular administration could also produce antitumor effects, we tested this approach using the same daily dose as the intraperitoneal treatments (Figure 1G). However, intramuscular administration did not provide any therapeutic benefit for PDAC tumors, as shown by IVIS imaging (Figure 1H). There was no significant difference in tumor bioluminescent intensity between the H Flu and PBS-treated groups (Figure 1I). These findings indicate that the antitumor effects of H Flu in orthotopic PDAC models require intraperitoneal administration.

### H Flu drives an effector phenotype in tumor-infiltrating lymphocytes in orthotopic PDAC

The sustained reduction in tumor progression even following the H Flu treatment suggests that its anticancer mechanism is at least in part immune-mediated. To identify key immunological contributors to the efficacy of H Flu, we conducted flow cytometry analysis of tumors from H Flu or PBS treated mice (Figure 2A). Paralleling our findings in the subcutaneous PDAC model, H Flu treatment also appeared to shift the TME from a suppressive state characterized by Tregs and MDSCs, to a T cell and NK cell, inflamed state (Figure 2B). Though a statistically significant increase in CD8+ and CD4+ T cell populations was not attained (Figure 2C), there was indeed an observable increase in these populations in H Flu treated mice. With respect to T cell functionality, H Flu treatment did significantly enhance activation of these T cell populations, measured by increased CD69 expression (Figure 2D). Further, intratumoral CD4+ T cells from H Flu treated mice exhibited enhanced cytotoxic functionality, measured by granzyme B and perforin expression (Figure 2E). Apart from T cells, increased infiltration of NK cells and reduced MDSC populations within the tumor are both associated with positive tumor outcomes^12,38^. In addition to enhanced T cell functionality, H Flu treated mice also experienced significant NK cell infiltration in the tumor (Figure 2F), while also reducing MDSC populations (Figure 2G). Together, these results parallel many of our initial findings in the subcutaneous model, in that H Flu treatment establishes an immunologically inflamed tumor state.

**Figure 2.**
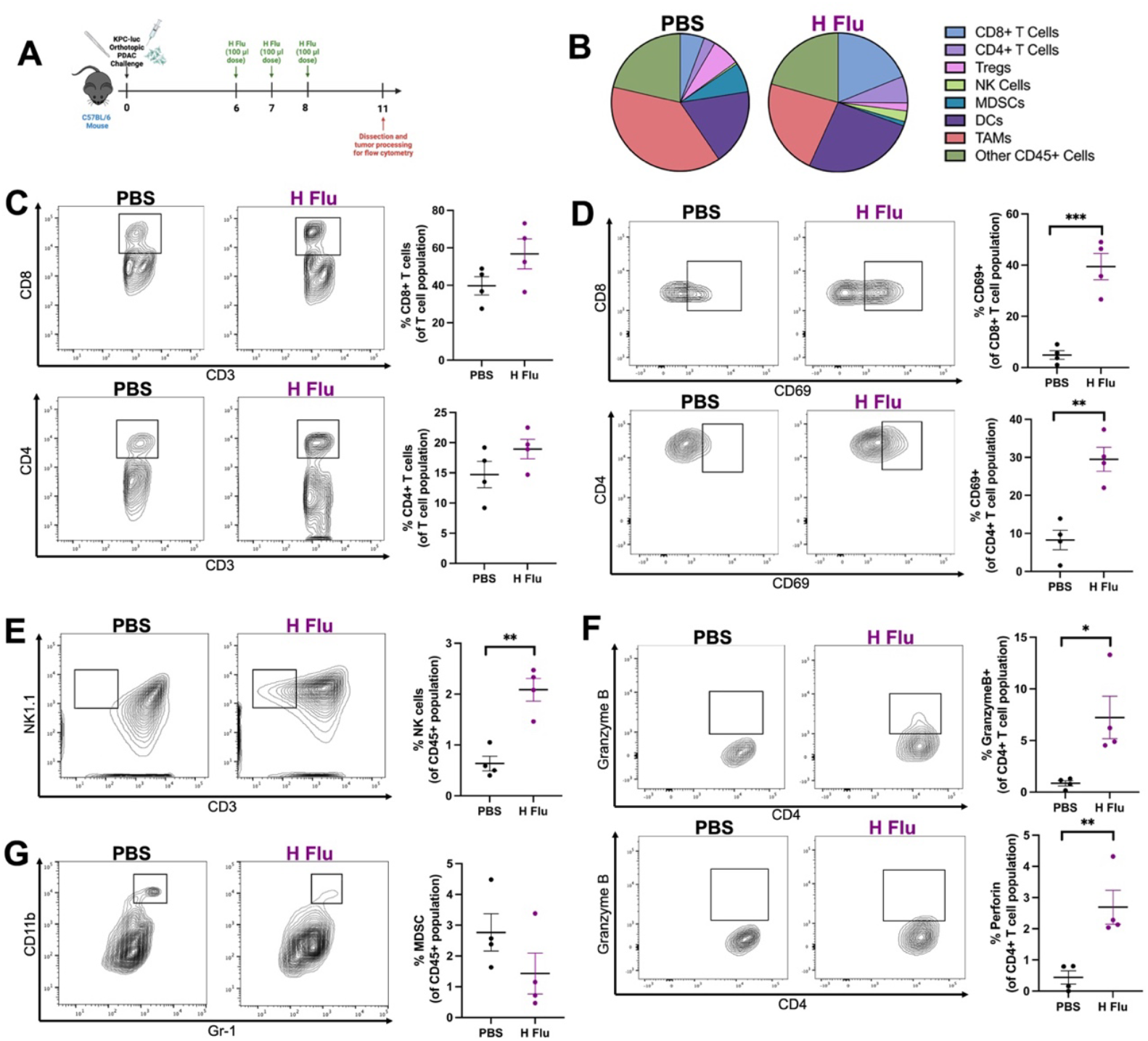
H Flu vaccine treatment reprograms PDAC tumor landscape in orthotopic model. (**A**) Experimental design describing dissection of KPC-luc tumors for flow cytometry analysis following completion of treatment with H Flu vaccine. n = 4 mice per group. (**B**) Representative pie charts of tumor-resident immune cell populations from experiment described in A. CD8+ T cells were defined as CD3+ CD4-CD8+, CD4+ T cells were defined as CD3+ CD4+ CD8-FoxP3-, regulatory T cells (Tregs) were defined as CD3+ CD4+ FOXP3+, natural killer (NK) cells were defined as NK1.1+ CD3-, myeloid derived suppressor cells (MDSCs) were defined as CD11b+ Gr1+, dendritic cells (DCs) were defined as CD11c+ MHCII+, tumor-associated macrophages (TAMs) were defined as CD11b+ F4/80+. (**C**) Representative flow cytometry plot of intratumoral CD8+ T cells (top left) and CD4+ T cells (bottom left), and graphs depicting CD8+ and CD4+ T cell populations for PBS and H Flu treated groups (top right and bottom right). (**D**) Representative flow cytometry plot of CD69-expressing CD8+ T (top left) and CD4+ T cell populations (bottom left), and graphs depicting CD69 expression for T cell populations from PBS and H Flu treated groups (top right and bottom right). (**E**) Representative flow cytometry plots of granzyme B (top left) and perforin (bottom left) expression from CD4+ T cells, and graphs depicting granzyme B (top right) and perforin (bottom right) expression for CD4+ T cells. (**F**) Representative flow cytometry plot of intratumoral NK cells (left) and graphs depicting NK cell populations from PBS and H Flu treated groups (right). (**G**) Representative flow cytometry plot of intratumoral MDSCs (left) and graphs depicting MDSC populations from PBS and H Flu treated groups (right) Two-tailed student’s T-test (C-E) *P < 0.05, **P < 0.01, ***P < 0.001.

### Pre-immunization with H. influenzae drives the development and accumulation of effector memory T cell populations within the tumor microenvironment

Memory-derived immune cells, particularly those specific to microbial antigens, have been shown to reactivate upon exposure. In cancer, these reactivated cells can be directed to achieve potent tumor eradication^16^. Thus, we investigated the effects immunizing mice with H Flu prior to tumor initiation, and then proceeded to attempt to reactivate immunological memory with H Flu once tumors are established (Figure 3A). Unexpectedly, just immunizing with H Flu prior to tumor challenge appeared to impede tumor development, resulting in smaller tumors compared to PBS-immunized mice (Figure 3B).

**Figure 3.**
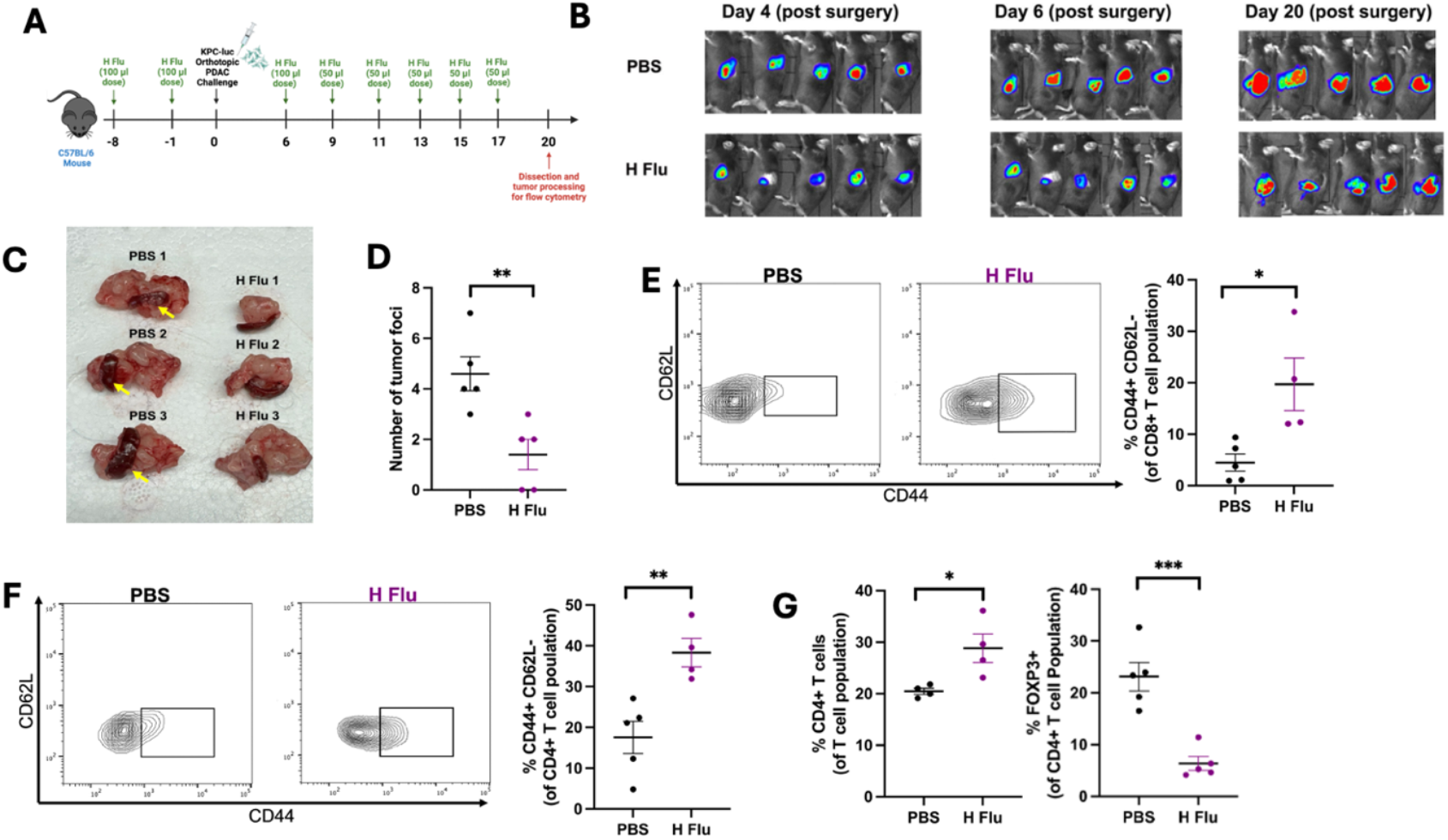
H Flu immunization establishes immunological memory and augments therapeutic efficacy. (**A**) Experimental design describing immunization and treatment schedule utilizing orthotopic PDAC tumor model. n = 4-5 mice per group. (**B**) Representative bioluminescent images of mice bearing KPC-luc tumors following H Flu or PBS treatment as described in panelA. (**C**) Examples of primary KPC-luc tumors and spleens following treatment schedule described in A. Yellow arrows denote PDAC metastatic foci localized to the spleen. (**D**) Graph depicting quantification of spleen metastasis following H Flu or PBS treatment. (**E**) Representative flow cytometry plot of intratumoral effector memory (CD44+ CD62L-) CD8+ T cells (left), and graph depicting effector memory CD8+ T cells for PBS and H Flu treated groups (right). (**F**) Representative flow cytometry plot of intratumoral effector memory (CD44+ CD62L-) CD4+ T cells (left), and graph depicting effector memory CD4+ T cells for PBS and H Flu treated groups (right). (**G**) Graphs depicting CD4+ and Treg (CD4+ FOXP3+) populations detected by flow cytometry of individual tumors for H Flu and PBS treated groups. Two-tailed student’s T-test (D-G). *P < 0.05, **P < 0.01, ***P < 0.001

In tumor-bearing mice, pre-immunization with H Flu further amplified the effects of the initial immunizations, leading to sustained tumor growth suppression throughout the duration of the study (Figure 3B). In addition to reducing primary tumor progression, H Flu immunization and re-exposure resulted in significant reduction in PDAC tumor foci present on the spleen (Figure 3C & 3D), suggesting this immunization schedule with also provides an antimetastatic effect in PDAC. To evaluate whether the physiological responses to H. influenzae were linked to the establishment of memory T cell populations, we performed flow cytometry on the dissected tumors. Strikingly, we observed nearly a 4-fold increase in effector memory CD8+ T cells and a 2-fold increase in effector memory CD4+ T cells within the tumors compared to the control group (Figure 3E & 3F). Additionally, H Flu -treated mice showed an overall rise in CD4+ T cells and a significant reduction in Treg populations, suggesting that effector memory T cells may contribute to the increased intratumoral CD4+ T cell population (Figure 3G). These findings highlight H. influenzae’s capacity to establish and recruit memory T cell populations to the tumor microenvironment effectively.

### Antitumor immunity from H Flu is mediated by CD4+ T cell and NK cell populations

Following H Flu treatment, tumors showed increased infiltration of NK cells, CD4+ T cells, and CD8+ T cells, prompting us to investigate the specific role of each population in the antitumor response. To determine the contribution of NK cells, we conducted an in vivo depletion using an anti-NK1.1 antibody (Figure 4A). The depletion of NK cells completely eliminated H Flu’s antitumor effect (Figure 4B, Supplemental Figures S2B and S3B), underscoring the essential role of NK cells in H Flu’s immunological mechanism.

**Figure 4.**
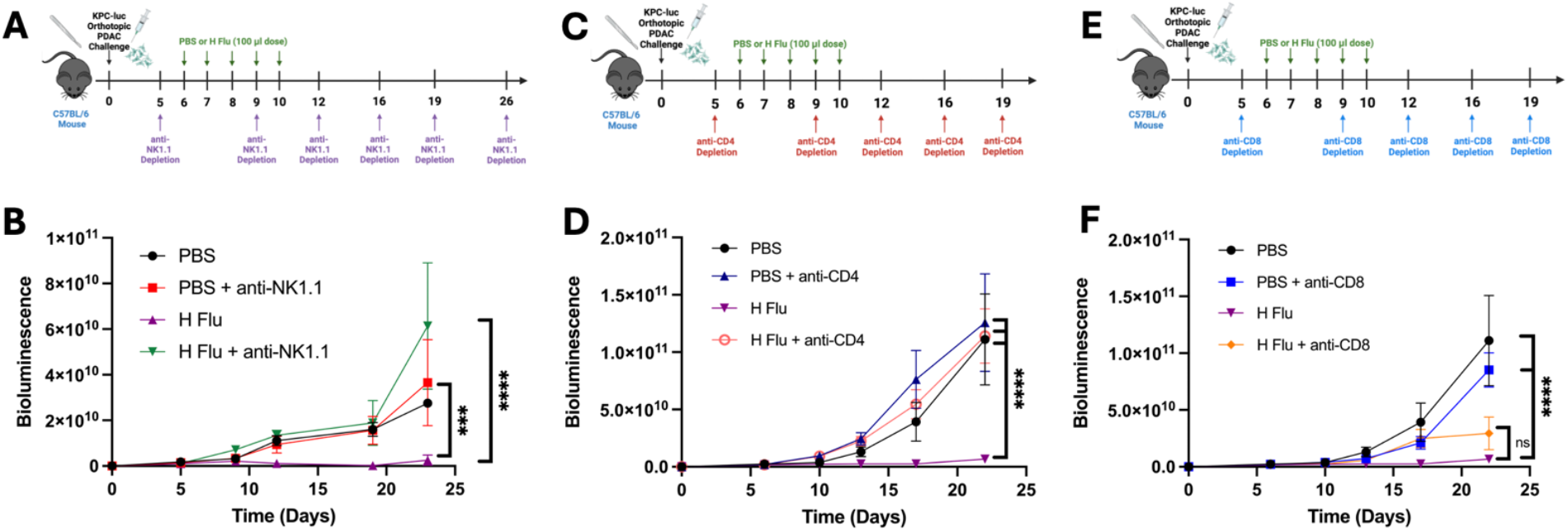
H Flu therapeutic efficacy is abrogated with depletion of CD4+ T cells and NK cells. (A) Experimental design describing H Flu treatment and NK cell depletion schedule utilizing an orthotopic PDAC tumor model. n = 3-5 mice per group. (B) Tumor growth curve plotting bioluminescent intensity (p-1 sec-1 cm2 sr-1) compared to time following the treatment regimen described in A. (C) Experimental design describing H Flu treatment and CD4+ T cell depletion schedule utilizing an orthotopic PDAC tumor model. n = 4-5 mice per group. (D) Tumor growth curve plotting bioluminescent intensity (p-1 sec-1 cm2 sr-1) compared to time following H Flu or PBS treatment and CD4+ T cell depletion. (E) Experimental design describing the H Flu treatment and CD8+ T cell depletion utilizing an orthotopic PDAC tumor model. n = 4-5 mice per group. (F) Tumor growth curve plotting bioluminescent intensity (p-1 sec-1 cm2 sr-1) compared to time following H Flu or PBS treatment and CD8+ T cell depletion. 2-way ANOVA with Tukey correction (C and G). ns – not significant, ***P < 0.001, ****P < 0.0001.

Given the marked reduction in H. influenzae’s antitumor response following NK cell depletion, we next performed in vivo depletion studies targeting CD4+ and CD8+ T cells individually (Figure 4C). IVIS imaging revealed substantial tumor progression in both H. influenzae- and PBS-treated groups within the CD4+ T cell-depleted cohort (Figure 5E), indicating that H. influenzae is unable to suppress tumor growth in the absence of CD4+ T cells. Consistently, the bioluminescent intensity and tumor size in H. influenzae-treated, CD4+ T cell-depleted mice were comparable to those in the PBS control group (Figure 4D, Supplemental Figures S2D and S3D). Notably, in CD8+ T cell-depleted mice treated with H. influenzae, the antitumor response was not completely eliminated (Figure 4D, Supplemental Figures S2D and S3D). However, the difference in tumor progression between H. influenzae-treated and CD8+ T cell-depleted groups was not statistically significant (Figure 4D). These findings indicate that the therapeutic effect of H. influenzae is primarily mediated by CD4+ T cells and NK cells, with a lesser role for CD8+ T cells.

**Figure 5.**
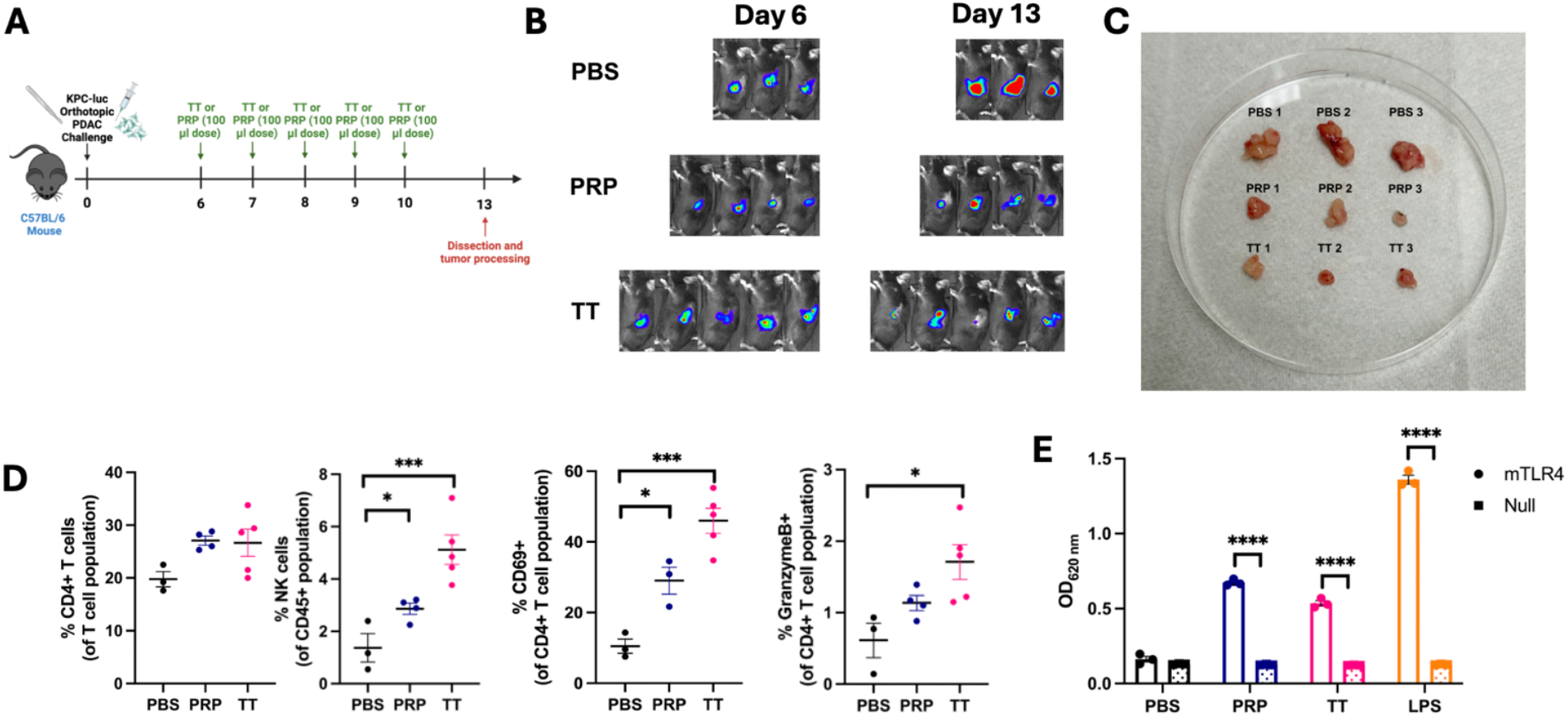
Individual H Flu Vaccine Components Elicit Antitumor Response in PDAC. (**A**) Experimental design describing H Flu vaccine component treatment schedule utilizing an orthotopic PDAC tumor model. n = 3-5 mice per group. (**B**) Representative bioluminescent images of mice bearing KPC-luc tumors following TT, PRP, or PBS treatment as described in A. (**C**) Examples of primary KPC-luc tumors following treatment schedule described in A. (**D**) Graphs depicting (top left) CD4+ T cell (CD3+ CD4+ CD8-),(top right) NK cell (NK1.1+ CD3-), and CD69+ CD4+ T cell populations detected by flow cytometry of individual tumors for TT, PRP, and PBS groups following treatment schedule described in A. **(E)** Stimulation of TLR4 expressing HEK cells and non-TLR expressing, parental (Null) cell lines following treatment with PBS (negative control), PRP (2 µg/mL), TT (2 µg/mL), or LPS (positive control, 10 ng/mL). 1-way ANOVA with Tukey’s multiple comparisons test (D), 2-way ANOVA with Sidak’s correction for multiple comparisons (F). **\***P < 0.05, **P < 0.01, ***P < 0.001, ****P < 0.0001.

### The tetanus toxoid component of H Flu contributes antitumor immunogenicity and direct interaction with PDAC cells

To better understand the mechanism underlying H Flu’s antitumor effects, we evaluated the individual contributions of its vaccine components. The Hiberix vaccine, which targets H Flu contains two key components: a tetanus toxoid (TT) conjugated to a polyribosyl-ribitol phosphate (PRP) component^37^. Using an orthotopic model of PDAC, tumor-bearing mice were treated with doses of tetanus toxoid (TT) or polyribosyl-ribitol phosphate (PRP) equivalent to those in the Hiberix vaccine (Figure 5A). IVIS imaging at the study’s conclusion demonstrated reduced tumor growth in both TT- and PRP-treated groups compared to the PBS control (Figure 5B). However, analysis of the dissected tumors revealed that TT-treated tumors were significantly smaller than those treated with PRP (Figure 5C), indicating that TT plays a more prominent role in the antitumor response induced by H. influenzae. Consistent with this, TT exhibited greater immunogenicity than PRP. While both components modestly increased CD4+ T cell infiltration into the tumor, they significantly promoted NK cell infiltration, with TT having the most pronounced effect (Figure 5D). Notably, TT treatment also enhanced the activation and cytotoxic function of CD4+ T cells, as evidenced by increased expression of CD69 and granzyme B, respectively (Figure 5D).

The innate immune system relies on pattern recognition receptors (PRRs) to detect microbial components and initiate immune responses. To investigate whether H Flu activates any specific PRR, we screened a range of Toll-like receptors (TLRs) and identified TLR4 as the primary receptor utilized by H. Flu (Figure S1I). Since both components of H Flu contribute to reducing PDAC tumor progression, with TT demonstrating greater cell-based immunogenicity, we next investigated their individual roles in activating TLR4. Interestingly, both TT and PRP significantly stimulated TLR4 activity compared to control Null cell lines (Figure 5E). These findings indicate that both components of H Flu contribute to its antitumor effects in PDAC by activating TLR4. However, TT appears to be the dominant contributor, likely due to its stronger immunogenic properties and ability to engage innate immune pathways more effectively.

### Tetanus toxoid mitigates PDAC aggressiveness independently of the immune system

Because TT elicited a strong antitumor effect than PRP, we next investigated potential tumor cell intrinsic mechanisms of TT, that are independent of its immunostimulatory qualities. Canonically, tetanus toxin binds to the sialic acid residues that comprise ganglioside GD3^39^. Because sialic acids have been heavily implicated in PDAC tumorigenesis^28^, we speculated that TT may be therapeutically targeting tumoral sialic acids. To explore the direct interaction between TT and PDAC cells, we performed an immunoprecipitation assay targeting TT (Figure 6A). The subsequent SDS-PAGE analysis of the immunoprecipitates revealed distinct protein bands in the TT and H Flu lanes that corresponded to the expected molecular weights of the heavy and light chains of TT. These bands were absent in the PBS control, confirming the specific interaction between TT and PDAC cells (Figure 6B).

**Figure 6.**
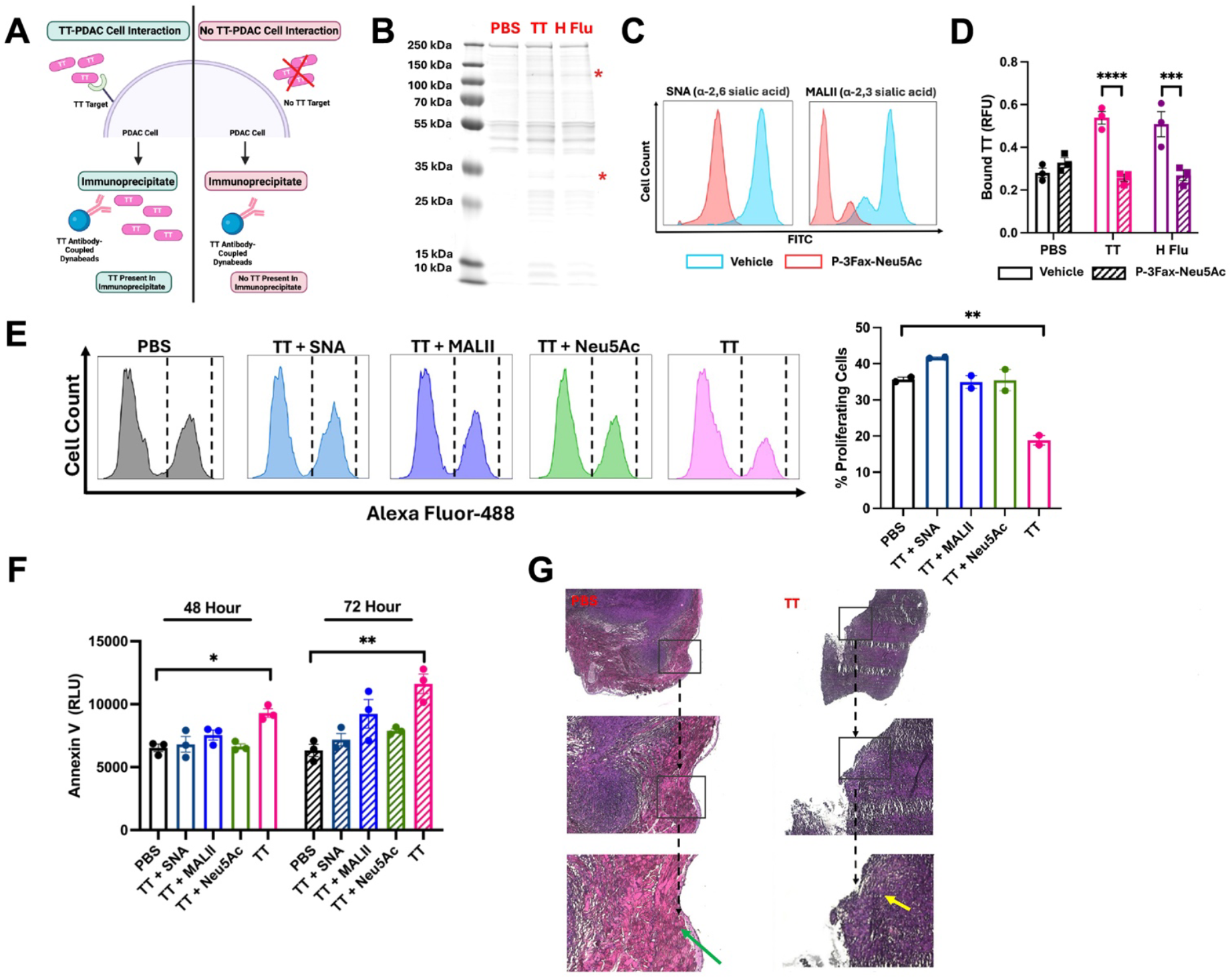
TT Interacts with Sialic Acids Expressed on PDAC Cells and Diminishes Tumor Cell Aggressiveness. (**A**) Schematic depicting immunoprecipitation of TT to identify direct interaction with 6419c5 PDAC cells. (**B**) Image of SDS-PAGE of TT-targeting immunoprecipitate from 6419c5 cells treated with PBS, TT, or H Flu. Red asterisks denote approximate molecular size of TT heavy (∼100 kDa) and light (∼50 kDa) chains. (**C**) Representative flow cytometry histograms depicting α-2,6 and and α-2,3 sialic acid expression on 6419c5 cells in the presence of vehicle (blue histogram) or sialyltransferase inhibitor P-3Fax-Neu5Ac (red histogram). (**D**) Measurement of bound tetanus toxoid on 6419c5 cells in the absence or presence of a sialyltransferase inhibitor. Cells were plated in triplicate for all treatment conditions. (**E**) Representative flow cytometry histograms obtained after treating PDAC cells with TT alone or combined with lectins or exogenous Neu5Ac. The dashed lines denote Click-iT EdU positive cell populations for each treatment (left). Measurement of proliferating cell populations (Click-iT positive) for each treatment (right). Samples were run in duplicate for all treatment conditions. (**F**) Effects of TT on apoptosis of 6419c5 cells alone or in combination with lectins or exogenous Neu5Ac. Each treatment was tested at 1 µg/mL and measured at different timepoints. (**G**) Representative images of H&E-stained sections from PBS or TT-treated tumors. Stained sections were imaged at 4X, 10X, and 20X, respectively. 1-way ANOVA with Tukey’s multiple comparisons test (E and F). 2-way ANOVA with Sidak’s correction for multiple comparisons (D). ***P < 0.001, ****P < 0.0001.

Next, we examined the expression of sialic acids on 6419c5 PDAC cells in the presence or absence of the sialyltransferase inhibitor P-3Fax-Neu5Ac (Figure 6C). To further investigate the potential role of sialic acids in the TT-PDAC cell interaction, we conducted a cell-based ELISA. We observed an increase in bound TT in both the H Flu and TT-treated groups, but the addition of P-3Fax-Neu5Ac resulted in a noticeable reduction in TT binding for both treatments (Figure 6D). Importantly, the PBS group showed no change in TT binding, even with reduced sialic acid expression, suggesting that the TT-sialic acid interaction was specifically affected by the inhibitor.

To examine the biological relevance of the TT-sialic acid interaction on PDAC cell behavior, we assessed its impact on cell proliferation using the Click-iT EdU assay. 6419c5 cells treated with TT exhibited a significant reduction in proliferation compared to PBS-treated cells (Figure 6E). To determine whether this effect was mediated through sialic acid binding, we tested the combination of TT with sialic acid-binding lectins Sambucus Nigra Lectin (SNA) and Maackia amurensis lectin (MALII) or exogenous sialic acid (Neu5Ac), aiming to compete for sialic acids or increase non-cell-based binding of TT. Interestingly, when combined with SNA, MALII, or Neu5Ac, the anti-proliferative effect of TT was diminished, with results comparable to the PBS-treated control (Figure 6E).

We further investigated the impact of TT on PDAC cell apoptosis using Annexin V staining. Treatment with TT led to increased apoptosis in PDAC cells, but this effect was completely abrogated when cells were co-treated with SNA, MALII, or Neu5Ac (Figure 6F), confirming the role of sialic acid in mediating TT-induced apoptosis.

Finally, to assess the effects of TT on tumor composition, we performed H&E staining on tumor sections from mice treated with either TT or PBS. Tumor sections from PBS-treated mice showed a prominent collagen-rich stromal layer surrounding the tumor periphery (Figure 6G, green arrow). In contrast, tumor sections from TT-treated mice lacked this dense stromal barrier and displayed increased cell infiltration within the tumor (Figure 6G, yellow arrows). These findings suggest that the direct interaction between TT and tumoral sialic acids may counteract the pro-tumorigenic effects of sialylation in PDAC, promoting a less supportive tumor microenvironment and enhancing tumor cell infiltration.

## Discussion

Pancreatic ductal adenocarcinoma remains a significant therapeutic challenge, particularly when it comes to immunotherapy. The disease’s inherently low immunogenicity, coupled with a highly immunosuppressive tumor microenvironment (TME) and minimal mutational burden, enables PDAC to evade detection by the immune system, making most ICI therapies largely ineffective. Immunotherapy and vaccination therapies aim to identify and target non-self antigens, which are present on tumor surfaces, by harnessing the body’s natural immune response^40,41^. These therapies have garnered increasing interest in the treatment of pancreatic ductal adenocarcinoma. Recent studies suggest that immunotherapies may offer advantages over traditional treatments, including reduced side effects and lower tumor resistance^42^. Microbial-based immunotherapies, in particular, hold promise as a novel approach by utilizing non-self-antigens to stimulate the immune system^43^. While these therapies show potential for both patients and the healthcare system, the field remains in its early stages. A deeper understanding of innovative therapeutic strategies for PDAC is urgently needed to realize their full potential^44^.

To overcome these limitations, we explored an innovative strategy by repurposing the FDA-approved Haemophilus influenzae serotype B (H Flu) vaccine as a potential immunotherapy for PDAC. This vaccine, originally developed for preventing infections, offered a safer, less toxic alternative to live bacterial therapies. In our study, we evaluated the anticancer efficacy of the H Flu vaccine using subcutaneous, orthotopic, and genetically engineered murine models of PDAC. Following the identification of the vaccine’s antitumor effects, we sought to investigate the underlying mechanisms driving its efficacy, focusing on both the immunological responses and changes in the tumor’s composition.

We show that systemic administration of H Flu is highly effective in reducing PDAC tumor progression and improving survival outcomes. This has a marked advantage over our previous studies, which necessitate intratumoral administration for an effective antitumor response, therefore the therapy is limited to superficial tumor types. Moreover, the H Flu antitumor response coincided with increases not only in tumor infiltration of CD4+, CD8+ T cells, and NK cell populations, but also significant increases cytotoxic or effector mediators such as CD69, granzyme B, and perforin. The critical function of CD4+ T cells and NK cells became apparent, when depletion of these populations resulted in complete attenuation of the H Flu antitumor response. The immunological mechanism of H Flu contrasts that of our previous studies investigating microbial-based vaccine immunotherapy, which were primarily CD8+ T cell mediated. Certainly, this difference may be attributed to the vaccines comprising components from different microbial sources, that have inherently preferential activation of immune cells.

In this study, we also demonstrate that the TT component of the H Flu vaccine plays a more significant role in driving antitumor immunity compared to the other component, PRP. This finding suggests that TT is likely the main contributor to the immunostimulatory effects of H Flu. Indeed, many of the immune populations activated by H Flu appear to overlap with those targeted by TT, further supporting its central role in mediating the immune response. This outcome is not entirely unexpected, as TT has previously been shown to function as an effective immune adjuvant in tumor antigen-based vaccine therapies, particularly for cancers such as breast and cervical cancer. By enhancing the immune response in these contexts, TT may offer similar benefits in the treatment of PDAC, reinforcing its potential as a valuable tool for cancer immunotherapy^45-47^.

This study sets itself apart from previous research on TT in cancer by going beyond its known immunostimulatory effects to explore a novel tumor-specific mechanism. We identified a previously unknown interaction between TT and tumor-associated sialic acids. While TT had been recognized for interacting with sialic acid residues on motor neurons, this study reveals a new role for TT in targeting sialic acids within tumors. Our findings show that this interaction helps counteract the anti-apoptotic and hyperproliferative effects of tumor cell sialylation. This newly discovered function of TT could be especially relevant for disrupting the stromal environment in PDAC, given the high levels of sialylation in its tumor stroma^48,49^. We observed a reduction in tumor stroma after TT treatment, which may be due to the interaction between TT and stromal sialic acids. While further investigation is needed, these results carry important therapeutic potential.

This research shares some similarities with studies on Listeria-mediated delivery of TT to PDAC tumors, though there are also notable differences^18^. It is not surprising that both studies involve TT as part of the treatment for PDAC. The most significant parallel between the two lies in the immunological mechanisms. Both H Flu and Listeria-TT were shown to enhance T cell effector function and promote CD4+ T cell-mediated, but not CD8+ T cell-mediated, antitumor immunity. Additionally, both treatments led to the formation of CD44+ CD62L-memory T cell populations following an immunization-reactivation treatment schedule^18^. However, there are key differences in the findings as well. In a survival study, Listeria-TT combined with gemcitabine extended survival of KPC mice by over 60 days compared to the control group, while the current research found that H Flu extended survival by nearly 160 days. One notable difference between the two approaches is that the H Flu vaccine contains the full, formaldehyde-inactivated tetanus toxin protein, while Listeria-TT uses a non-toxic fragment of the protein. This difference suggests that the TT in H Flu may be more immunogenic, potentially explaining the greater survival benefit observed in this study. Additionally, the requirement for NK cells in H Flu’s antitumor immunity, which was not reported in the Listeria-TT study, may be linked to the presence of the whole TT protein in H Flu.

A significant portion of this study was conducted in an orthotopic PDAC model using the KPC-luc cell line to develop tumors. Recently, KPC-luc tumors were identified to possess higher baseline proportions of CD8+ T cells and NK cells residing within the tumor compared to non-luciferase expressing, KPC tumors. Importantly, KPC-luc tumors were also significantly smaller than KPC tumors implanted the same day^50^. The findings of this study are relevant to our study because the immune cell populations we observed H Flu to enrich are already present within the tumor, according to this study. Further, the higher proportions of these effector cell populations are also not reflective of clinical PDAC. With this consideration, it would be prudent to conduct subsequent studies in the KPC model, or a non-luciferase expressing orthotopic model of PDAC.

In summary, this study demonstrates identifies a novel dual-faceted role of tetanus toxoid as an anticancer agent that also significantly contributes to the antitumor responses we observed from the H Flu vaccine. H Flu treatment is effective in directing immunological memory cells to PDAC tumors, as well as mitigating tumor cell aggression through sialic acid targeting. We propose H Flu and TT as promising, dual-faceted therapeutics to be utilized against PDAC.

## Authors Contribution

**E.G**. Contributed to data curation, formal analysis, validation, visualization, methodology development, project administration, and the initial drafting of the manuscript, review and editing

**S.P**. Concept, funding acquisition, resources, and reviewed the manuscript.

**K.G**. Conceptualization and design of the study, data curation and investigation, resources, visualization, methodology development, supervision, project administration, and manuscript writing, review, and editing.

## References

1. Bray F, Laversanne M, Sung H, et al. Global cancer statistics 2022: GLOBOCAN estimates of incidence and mortality worldwide for 36 cancers in 185 countries. CA Cancer J Clin. 2024; 74(3): 229–263.

2. Siegel, Rebecca L., et al. “Cancer statistics, 2023.” CA: a cancer journal for clinicians 73.1 (2023).

3. Bear, Adham S., Robert H. Vonderheide, and Mark H. O’Hara. “Challenges and opportunities for pancreatic cancer immunotherapy.” Cancer cell 38.6 (2020): 788–802.

4. Principe, Daniel R., et al. “Trials and tribulations of pancreatic cancer immunotherapy.” Cancer letters 504 (2021): 1–14.

5. Giurini, Eileena F., et al. “Looking Beyond Checkpoint Inhibitor Monotherapy: Uncovering New Frontiers for Pancreatic Cancer Immunotherapy.” Molecular Cancer Therapeutics (2024): OF1–OF15.

6. Waldman, Alex D., Jill M. Fritz, and Michael J. Lenardo. “A guide to cancer immunotherapy: from T cell basic science to clinical practice.” Nature Reviews Immunology 20.11 (2020): 651–668.

7. Guo, Mengdi, Melissa Yi Ran Liu, and David G. Brooks. “Regulation and impact of tumor-specific CD4+ T cells in cancer and immunotherapy.” Trends in Immunology (2024).

8. Page, Audrey, et al. “Development of NK cell-based cancer immunotherapies through receptor engineering.” Cellular & Molecular Immunology 21.4 (2024): 315–331.

9. Borst, Jannie, et al. “CD4+ T cell help in cancer immunology and immunotherapy.” Nature Reviews Immunology 18.10 (2018): 635–647.

10. Tay, Rong En, Emma K. Richardson, and Han Chong Toh. “Revisiting the role of CD4+ T cells in cancer immunotherapy—new insights into old paradigms.” Cancer gene therapy 28.1 (2021): 5–17.

11. Bawden, Emma G., et al. “CD4+ T cell immunity against cutaneous melanoma encompasses multifaceted MHC II–dependent responses.” Science immunology 9.91 (2024): eadi9517.

12. Chu, Junfeng, et al. “Natural killer cells: a promising immunotherapy for cancer.” Journal of translational medicine 20.1 (2022): 240.

13. Gupta, Kajal H., et al. “Bacterial-based cancer therapy (BBCT): recent advances, current challenges, and future prospects for cancer immunotherapy.” Vaccines 9.12 (2021): 1497.

14. Diwan, Deepti, et al. “Microbial cancer therapeutics: A promising approach.” Seminars in cancer biology. Vol. 86. Academic Press, 2022.

15. Nguyen, Dinh-Huy, et al. “Bioengineering of bacteria for cancer immunotherapy.” nature communications 14.1 (2023): 3553.

16. Nouri, Yasmin, Robert Weinkove, and Rachel Perret. “T-cell intrinsic Toll-like receptor signaling: implications for cancer immunotherapy and CAR T-cells.” Journal for ImmunoTherapy of Cancer 9.11 (2021).

17. Duong, Mai Thi-Quynh, et al. “Bacteria-cancer interactions: bacteria-based cancer therapy.” Experimental & molecular medicine 51.12 (2019): 1–15.

18. Selvanesan, Benson Chellakkan, et al. “Listeria delivers tetanus toxoid protein to pancreatic tumors and induces cancer cell death in mice.” Science translational medicine 14.637 (2022): eabc1600.

19. Raman, Vishnu, et al. “Intracellular Salmonella delivery of an exogenous immunization antigen refocuses CD8 T cells against cancer cells, eliminates pancreatic tumors and forms antitumor immunity.” Frontiers in Immunology 14 (2023): 1228532.

20. Newman, Jenna H., et al. “Intratumoral injection of the seasonal flu shot converts immunologically cold tumors to hot and serves as an immunotherapy for cancer.” Proceedings of the National Academy of Sciences 117.2 (2020): 1119–1128.

21. Gupta, Kajal H., Eileena F. Giurini, and Andrew Zloza. “Seasonal influenza vaccines differentially activate and modulate toll-like receptor expression within the tumor microenvironment.” Frontiers in Oncology 14 (2024): 1308651.

22. Giurini, Eileena F., et al. “Inactivated sars-cov-2 reprograms the tumor immune microenvironment and improves murine cancer outcomes.” bioRxiv (2022): 2022–06.

23. Daniels, Preston, et al. “Intratumoral influenza vaccine administration attenuates breast cancer growth and restructures the tumor microenvironment through sialic acid binding of vaccine hemagglutinin.” International journal of molecular sciences 25.1 (2023): 225.

24. Büll, Christian, et al. “Sialic acids sweeten a tumor’s life.” Cancer research 74.12 (2014): 3199–3204.

25. Nakamori, Shoji, et al. “Increased expression of sialyl Lewisx antigen correlates with poor survival in patients with colorectal carcinoma: clinicopathological and immunohistochemical study.” Cancer research 53.15 (1993): 3632–3637.

26. Hodgson, Kirsty, et al. “Sialic acid blockade inhibits the metastatic spread of prostate cancer to bone.” EBioMedicine 104 (2024).

27. Lumibao, Jan C., et al. “Altered glycosylation in pancreatic cancer and beyond.” Journal of Experimental Medicine 219.6 (2022): e20211505.

28. Dobie, Christopher, and Danielle Skropeta. “Insights into the role of sialylation in cancer progression and metastasis.” British Journal of Cancer 124.1 (2021): 76–90.

29. Rodriguez, Ernesto, et al. “Sialic acids in pancreatic cancer cells drive tumour-associated macrophage differentiation via the Siglec receptors Siglec-7 and Siglec-9.” Nature communications 12.1 (2021): 1270.

30. Boelaars, Kelly, and Yvette van Kooyk. “Targeting myeloid cells for cancer immunotherapy: Siglec-7/9/10/15 and their ligands.” Trends in Cancer (2024).

31. Li, Jinyang, et al. “Tumor cell-intrinsic factors underlie heterogeneity of immune cell infiltration and response to immunotherapy.” Immunity 49.1 (2018): 178–193.

32. Lu, Shao-Wei, et al. “IL-20 antagonist suppresses PD-L1 expression and prolongs survival in pancreatic cancer models.” Nature communications 11.1 (2020): 4611.

33. Lagundžin, Dragana, et al. “An optimized co-immunoprecipitation protocol for the analysis of endogenous protein-protein interactions in cell lines using mass spectrometry.” STAR protocols 3.1 (2022): 101234.

34. Bonanni, Paolo, et al. “Recommended vaccinations for asplenic and hyposplenic adult patients.” Human vaccines & immunotherapeutics 13.2 (2017): 359–368.

35. Giurini, Eileena F., et al. “Microbial-derived toll-like receptor agonism in cancer treatment and progression.” Cancers 14.12 (2022): 2923.

36. Mallya, Kavita, et al. “Modeling pancreatic cancer in mice for experimental therapeutics.” Biochimica et Biophysica Acta (BBA)-Reviews on Cancer 1876.1 (2021): 188554.

37. GlaxoSmithKline. Hiberix (2024). U.S. Food and Drug Administration. Available online at: https://www.fda.gov/media/77017/download?attachment (Accessed May 2024)

38. Ager, Casey R., et al. “High potency STING agonists engage unique myeloid pathways to reverse pancreatic cancer immune privilege.” Journal for immunotherapy of cancer 9.8 (2021).

39. Chen, Chen, et al. “Gangliosides as high affinity receptors for tetanus neurotoxin.” Journal of Biological Chemistry 284.39 (2009): 26569–26577.

40. Balachandran, V. P., Beatty, G. L., & Dougan, S. K. (2019). Broadening the impact of immunotherapy to pancreatic cancer: challenges and opportunities. Gastroenterology, 156(7), 2056–2072.

41. Bockorny, Bruno, Joseph E. Grossman, and Manuel Hidalgo. “Facts and hopes in immunotherapy of pancreatic cancer.” Clinical Cancer Research 28.21 (2022): 4606–4617.

42. Bailey, P., Chang, D. K., Forget, M. A., Lucas, F. A. S., Alvarez, H. A., Haymaker, C., … & Roszik, J. (2016). Exploiting the neoantigen landscape for immunotherapy of pancreatic ductal adenocarcinoma. Scientific reports, 6(1), 35848.

43. Chandra, V., & McAllister, F. (2021). Therapeutic potential of microbial modulation in pancreatic cancer. Gut, 70(8), 1419–1425.

44. Sally, Á., McGowan, R., Finn, K., & Moran, B. M. (2022). Current and future therapies for pancreatic ductal adenocarcinoma. Cancers, 14(10), 2417.

45. Stergiou, Natascha, et al. “Reduced breast tumor growth after immunization with a tumor-restricted MUC1 glycopeptide conjugated to tetanus toxoid.” Cancer Immunology Research 7.1 (2019): 113–122.

46. Laubreton, Daphné, et al. “The fully synthetic MAG-Tn3 therapeutic vaccine containing the tetanus toxoid-derived TT830-844 universal epitope provides anti-tumor immunity.” Cancer Immunology, Immunotherapy 65 (2016): 315–325.

47. Alson, Donia, et al. “Combination vaccination with tetanus toxoid and enhanced tumor-cell based vaccine against cervical cancer in a mouse model.” Frontiers in Immunology 11 (2020): 927.

48. Egan, Hannah, et al. “Targeting stromal cell sialylation reverses T cell-mediated immunosuppression in the tumor microenvironment.” Cell Reports 42.5 (2023).

49. Boelaars, Kelly, et al. “Pancreatic cancer-associated fibroblasts modulate macrophage differentiation via sialic acid-Siglec interactions.” Communications Biology 7.1 (2024): 430.

50. Ferrari, Daniele Pereira, et al. “KPC-luciferase-expressing cells elicit an anti-tumor immune response in a mouse model of pancreatic cancer.” Scientific Reports 14.1 (2024): 13602.

