## Supplemental data 1 for "Tetanus Toxoid Utilizes Dual-Faceted Anticancer Mechanism Through Targeting Tumoral Sialic Acids and Enhancing Cytotoxic CD4+ T cell Responses Against Pancreatic Cancer"

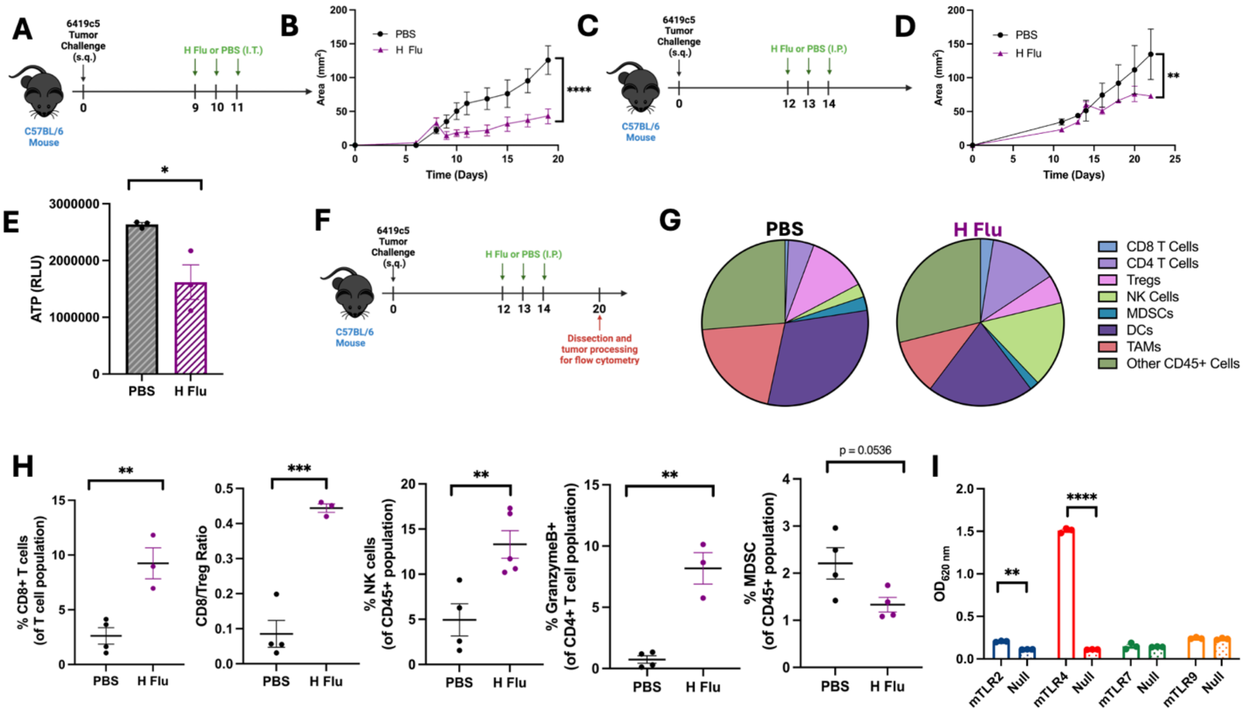


**Supplemental Figure S1:** Screening of Microbial-Based Vaccines for Antitumor Properties in PDAC. (**A**) Experimental design describing intratumoral treatment schedule with vaccines. n = 5 mice per group. (**B**) Tumor growth curves of 6419c5 tumors following treatment described in A. (**C**) Experimental design describing systemic (intraperitoneal) treatment schedule with vaccines. n = 3-5 mice per group. (**D**) Tumor growth curves of 6419c5 tumors following treatment described in C. (**E**) Measurement of metabolically active 6419c5 cells following PBS or H Flu treatment. Cell lines were plated in triplicate for experiment. (**F**) Experimental design describing dissection of 6419c5 tumors for flow cytometry analysis following completion of treatment with H Flu vaccine. n = 3-5 mice per group. (**G**) Representative pie charts of tumor-resident immune cell populations from experiment described in F. CD8+ T cells were defined as CD3+ CD4- CD8+, CD4+ T cells were defined as CD3+ CD4+ CD8- FoxP3-, regulatory T cells (Tregs) were defined as CD3+ CD4+ FOXP3+, natural killer (NK) cells were defined as NK1.1+ CD3-, myeloid derived suppressor cells (MDSCs) were defined as CD11b+ Gr1+, dendritic cells (DCs) were defined as CD11c+ MHCII+, tumor-associated macrophages (TAMs) were defined as CD11b+ F4/80+. (**H**) Graphs depicting immune cell populations detected by flow cytometry of individual tumors for H Flu and PBS groups from the experiment described in F. (**I**) Stimulation of TLR expressing HEK cells and non-TLR expressing, parental (Null) cell lines following H Flu stimulation. Cell lines were plated in triplicate for experiment. 2-way ANOVA with Sidak's correction (B and D), two-tailed student's T-test (E and H), 2-way ANOVA with Sidak's correction for multiple comparisons (I). $^{*}P < 0.05$, $^{**}P < 0.01$, $^{***}P < 0.001$, $^{****}P < 0.0001$.

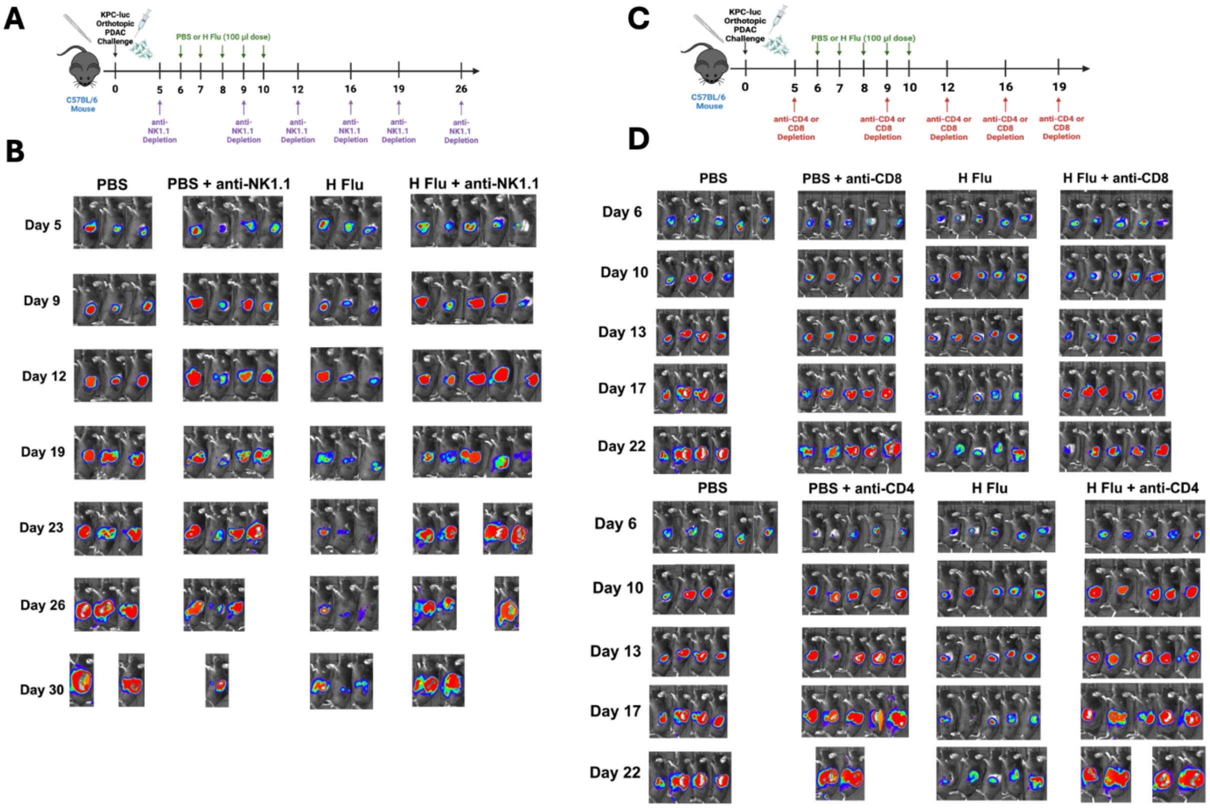


**Supplemental Figure S2**: Reduced tumor burden from H Flu treatment is mediated by CD4+ T cells and NK cells. (**A**) Experimental design describing H Flu treatment and NK cell depletion schedule utilizing an orthotopic murine PDAC model. n = 3-5 mice per group. (**B**) Representative bioluminescent images of mice bearing KPC-luc tumors following H Flu or PBS treatment and NK cell depletion as described in A, in which bioluminescent intensity is measured with radiance (p$^{-1}$ sec$^{-1}$ cm$^{2}$ sr$^{-1}$). (**C**) Experimental design describing H Flu treatment and CD4+ or CD8+ T cell depletion schedule utilizing an orthotopic PDAC tumor model. n = 4-5 mice per group. (**D**) Representative bioluminescent images of mice bearing KPC-luc tumors following H Flu or PBS treatment and CD8+ or CD4+ T cell cell depletion as described in C, in which bioluminescent intensity is measured with radiance (p$^{-1}$ sec$^{-1}$ cm$^{2}$ sr$^{-1}$).

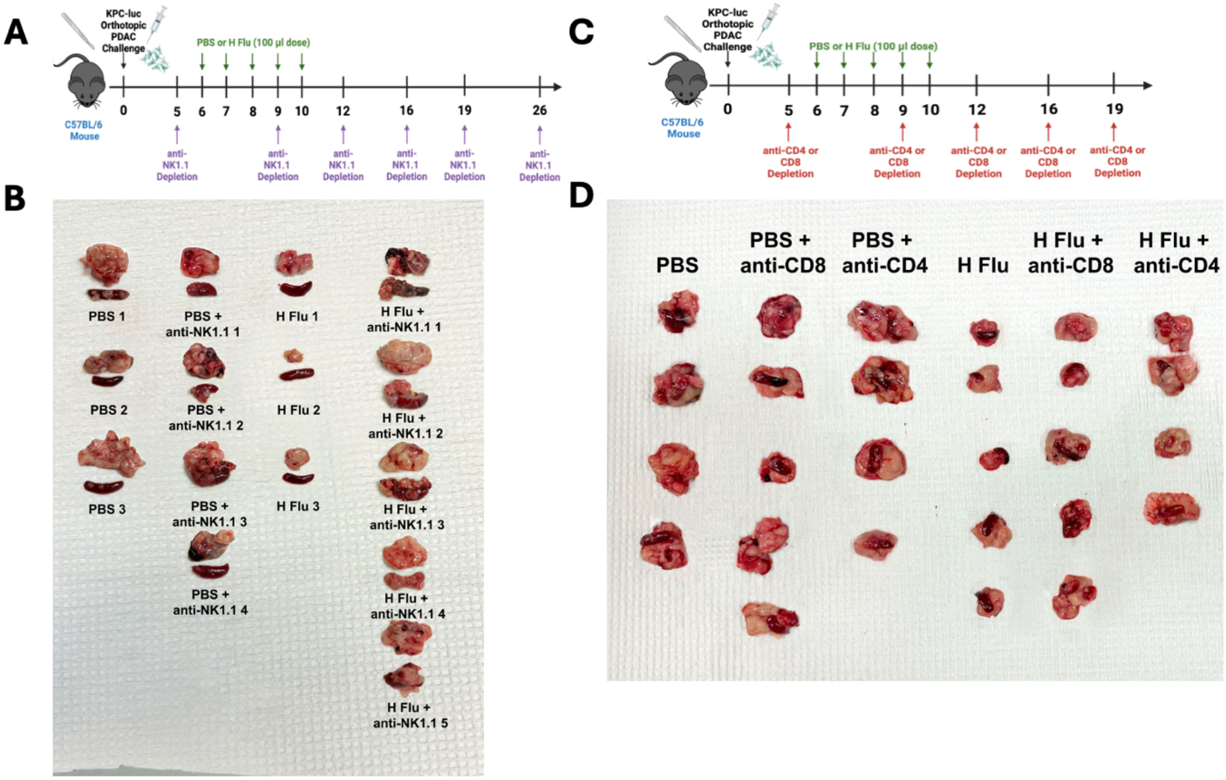


**Supplemental Figure S3:** Depletion of CD4+ T cells and NK cells arrests antitumor response from H Flu. (**A**) Experimental design describing H Flu treatment and NK cell depletion schedule utilizing an orthotopic PDAC tumor model. n = 3-5 mice per group. (**B**) Examples of primary KPC-luc tumors and spleens at study endpoint following in A. (**C**) Experimental design describing H Flu treatment and CD4+ or CD8+ T cell depletion schedule utilizing an orthotopic PDAC tumor model. n = 4-5 mice per group. (**D**) Examples of primary KPC-luc tumors and spleens at study endpoint following depletion schedule described in C.

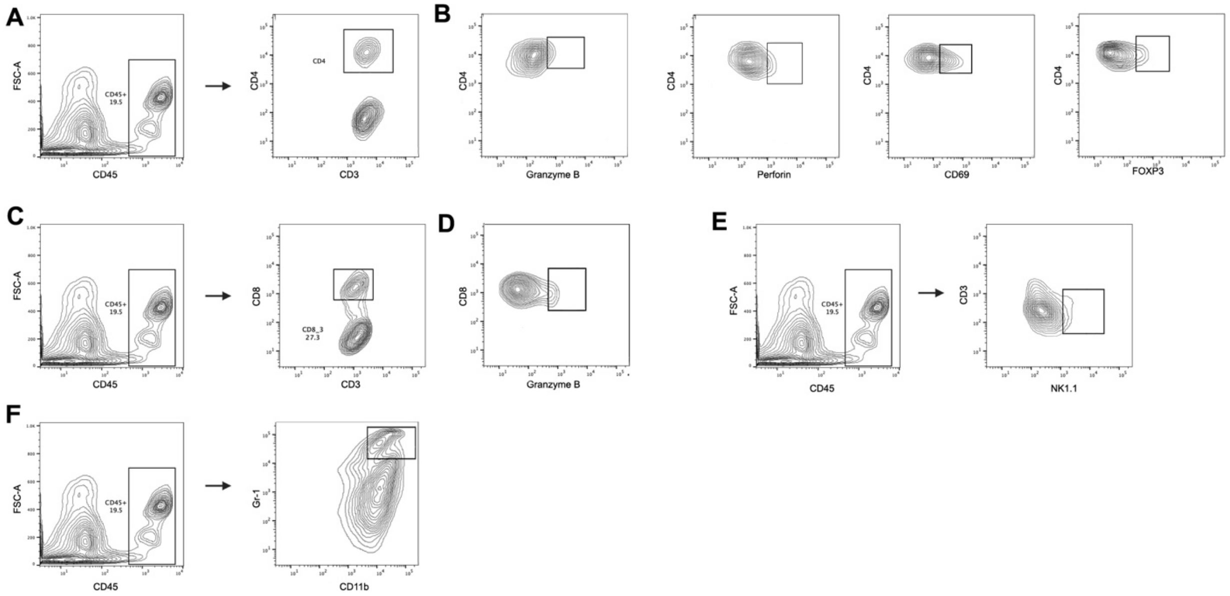


**Supplemental Figure S4:** Flow cytometry gating strategy of tumor resident immune cell populations. (**A)** Representative gating of CD4+ T cells from the CD45+ cell population. The CD4+ T cell population was defined as CD3+ CD4+. (**B**) Representative gating of perforin, CD69, and FOXP3 expressed on CD4+ T cells. (**C**) Representative gating of CD8+ T cells from CD45+ cell population. (**D)** Representative gating of granzyme B expression on CD8+ T cells. The CD8+ T cell population was defined as CD3+ CD8+. (**E**) Representative gating of NK cells from the CD45+ cell population. The NK cell population was defined as CD3- NK1.1+. (**F**) Representative gating of MDSCs from the CD45+ cell population. The MDSC population was defined as CD11b+ Gr-1+.

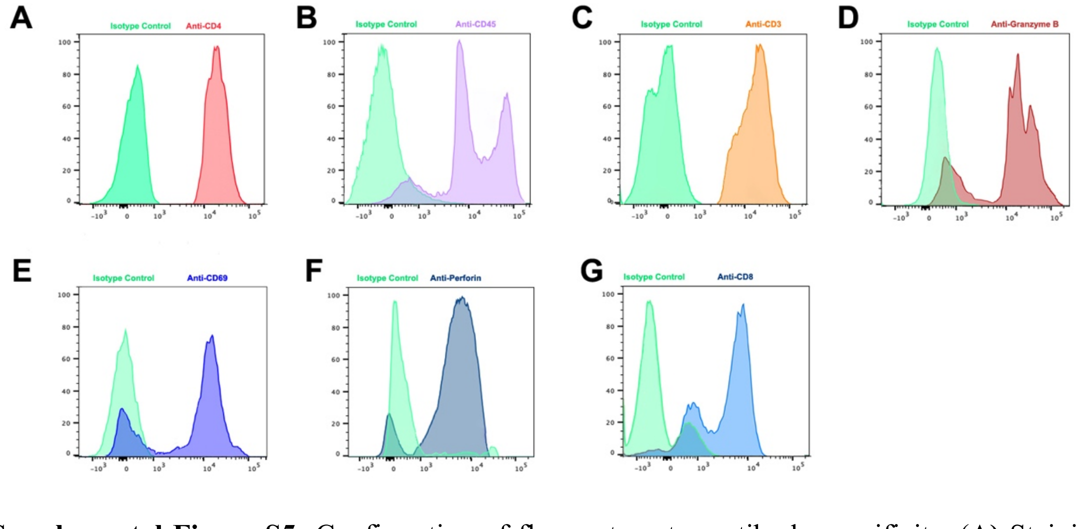


**Supplemental Figure S5:** Confirmation of flow cytometry antibody specificity. (**A)** Staining of CD45+ cells with CD4 targeting antibody or isotype control. (**B**) Staining with CD45 targeting antibody or isotype control. (**C**) Staining of CD45+ cells with CD3 targeting antibody or isotype control. (**D**) Staining of CD4+ T cells with granzyme B targeting antibody or isotype control. (**E**) Staining of CD4+ T cells with CD69 targeting antibody or isotype control. (**F**) Staining of CD4+ T cells with perforin targeting antibody or isotype control. (**G**) Staining of CD45+ cells with CD8 targeting antibody or isotype control.
